# A computational model of task allocation in social insects – ecology and interactions alone can drive specialisation

**DOI:** 10.1101/315846

**Authors:** Rui Chen, Bernd Meyer, Julian García

## Abstract

Social insect colonies are capable of allocating their workforce in a decentralised fashion; addressing a variety of tasks and responding effectively to changes in the environment. This process is fundamental to their ecological success, but the mechanisms behind it remain poorly understood. While most models focus on internal and individual factors, empirical evidence highlights the importance of ecology and social interactions. To address this gap we propose a game theoretical model of task allocation. Individuals are characterised by a trait that determines how they split their energy between two prototypical tasks: foraging and regulation. To be viable, a colony needs to learn to adequately allocate its workforce between these two tasks. We study two different processes: individuals can learn relying exclusively on their own experience, or by using the experiences of others via social learning. We find that social organisation can be determined by the ecology alone, irrespective of interaction details. Weakly specialised colonies in which all individuals tend to both tasks emerge when foraging is cheap; harsher environments, on the other hand, lead to strongly specialised colonies in which each individual fully engages in a single task. We compare the outcomes of self-organised task allocation with optimal group performance. Counter to intuition, strongly specialised colonies perform suboptimally, whereas the group performance of weakly specialised colonies is closer to optimal. Social interactions lead to important differences when the colony deals with dynamic environments. Colonies whose individuals rely on their own experience are more flexible when dealing with change. Our computational model is aligned with mathematical predictions in tractable limits. This different kind of model is useful in framing relevant and important empirical questions, where ecology and interactions are key elements of hypotheses and predictions.

## Introduction

Social insects are among the ecologically most successful life forms. They live in elaborately organised colonies, capable of managing a complex network of simultaneous tasks; from scouting and foraging to colony defence, nest building, thermoregulation, and brood care. One of the key factors for their ecological success is the colony’s ability to efficiently allocate its workforce to these different tasks, responding to frequent changes in external conditions and internal requirements [9, 35, 40, 62, 39, 10, 19, 22, 27, 30, 51, 59, 70]. The resulting “division of labour” (DOL) emerges from the task choices of individual workers without any central coordination or control. Deciphering the individual-based rules behind task selection is thus at the heart of understanding how colonies can achieve their collective plasticity in task allocation.

Task allocation in social insects has received a significant amount of attention [2]. The majority of research has focused on the influence of internal factors such as genetics [61], morphology [62] and hormones [69]. Comparatively, little attention has been given to the underlying mechanisms of social interactions and their role in regulating task allocation. This is in part because the dominant analysis framework does not provide a direct link between interactions among individuals and environmental conditions. Investigating the mechanistic roles of these factors has recently been proposed as the foundation to a more comprehensive understanding of DOL [30].

Mathematical and computational models have played an important role in this discussion – see [3, 19] for comprehensive reviews. Among these, response threshold models are arguably the most established [47, 43]. These models assume individuals have an internal task-related response threshold, and they are more likely to respond to a task stimulus the more it exceeds this threshold [4, 77, 63, 45, 32, 33, 18]. This simple idea has been implemented in many different forms, but regardless of the modelling details, interactions among individuals are always indirect. They only take place through stigmergy: individual actions change the task-related stimulus (for example, nest temperature for a task of thermoregulation) and the modified stimulus can in turn influence the task readiness of other individuals. Direct interactions do not feature in the models. This is in contrast to the empirical literature, which suggests that so-cial context is a significant determinant of task engagement [15, 31, 34].

In addition, the idea of stimulus intensity does not capture enough information to account for complex ecological factors. The cost required to tackle a task as well as the reward obtained from performing it can vary widely depending on environmental conditions. For example, how costly and successful a foraging task is can depend on factors such as ambient temperature, abundance of food sources and presence of predators. This in turn should be expected to affect the individuals’ decisions about task engagement. In other words, response threshold models do not provide a sufficient link to some empirically well-established environmental factors.

To overcome these limitations we propose a new modelling framework based on game theory. It directly incorporates social interactions and environmental conditions as the main factors. Although game theory is a well-established toolbox for the study of interdependent decision-making and has been very successfully applied to many other aspects of sociality in biology [42, 58, 20, 6], it has gone virtually unnoticed in the study of task allocation. An exception is [81], which investigates DOL in co-viruses. However, this study addresses evolutionary timescales rather than behavioural change in an individual’s lifetime.

We use Evolutionary Game Theory (EGT) to investigate how individual task preferences develop in the lifetime of a colony, and how specialisation can emerge as a result of these choices [57, 7]. It is important to note that we do not make any reference to evolutionary processes. Rather, our model addresses behavioural change in the scale of a colony lifetime. We are not the first to use EGT in this way. EGT has been used effectively to model behavioural change in ecological time-scales [42, 80].

We model task allocation as a simple game. Individuals choose how to divide their energy between different tasks. Task performance results in rewards that can be shared between colony members or go directly to the individual. Rewards are modulated by collective levels of investment into the tasks as well as by environmental factors. Groups of individuals repeatedly engage in collective task execution and modify their strategies based on the rewards they receive individually or collectively. This framework is aligned with empirical evidence identifying social interactions and ecology as key factors [24, 66, 68, 44, 31, 34, 15]. The framework provides a large degree of flexibility via simulations, while also allowing for mathematical predictions based on game theory [38].

The proposed formal model of task allocation directly addresses the influence of environmental factors and social interactions. Our model shows that ecological conditions are a crucial determinant for the emergence of different forms of specialisation. We also find that specialisation can emerge from interactions between individuals alone, under a large range of environmental conditions. Contrary to intuition, we find that specialisation does not always result in colony efficiency. Our theoretical results thus point towards promising new directions for empirical work that can bring us a step closer to understanding how social insects achieve their outstanding ecological success.

## Models and methods

### A task-allocation game

Any colony needs to balance its workforce allocation between different tasks in response to the conditions of the environment. As there is no centralised control mechanism, this allocation can only emerge from the task engagement decisions that individuals make. From the perspective of the individual, task engagement can be driven by the costs and benefits of the task and by the task choices of other individuals. This suggests game theory as an appropriate framework to investigate task allocation mechanisms. More specifically, shared benefits and individual costs place our scenario squarely into the context of public goods games [74, 1]. While we use the term “benefits to the individual”, in social insects this may well be a proxy for colony benefit rather than a direct benefit to the individual. As an example, individual workers can estimate the level of hunger in the colony by the rate of contact with hungry individuals. A reduction in this rate could constitute a benefit perceived by an individual.

We assume a large group of workers in a colony, who need to balance two prototypical tasks with different characteristics. Costs and benefits for each task vary under different ecological conditions. The benefits of task execution are shared by the whole colony, whereas costs accrue to individual workers. The payoff to a single individual equals the shared benefits generated by the task performance of the whole colony minus the individual cost of performing the tasks. Payoffs can be thought of as primarily metabolic or energetic in nature.

We use EGT to investigate how individuals modify and adjust their task choice behaviour based on simple rules that take only individual experience and social information into account. Importantly, unlike in classical game theory, there is no underlying assumption of rationality for the individuals and the processes are not driven by striving for collective efficiency or optimality. Only individual behaviour enters explicitly into the model, and colony-level task allocation patterns arise as an emergent property.

In our model individuals can choose between two tasks: a regulatory task *T* (for thermo-regulation) and a maximising task *F* (for foraging). *T* represents a homeostatic task: colonies need to maintain nest and brood temperature within certain bounds, for which a certain amount of collective effort is required. Allocating too little collective effort to this task can lead to regulation failure, and allocating too much effort does not improve the homeostasis and may in fact lead to suboptimal regulation, such as overcooling. Thus, group benefit for *T* as a function of group effort is a strictly concave function: the maximum benefit is obtained at an intermediate level of group effort. On the other hand, foraging is a maximising task; i.e., the benefit from *F* is monotonically increasing with the collective effort devoted to foraging: the more food is collected, the better.

Individual task preferences are determined by an inherent response trait [21, 46, 29]. The response trait of an individual *i* is modelled as a continuous value 0 ≤ *x*_*i*_ ≤ 1, which represents the fraction of effort that individual *i* invests into task *T*. Conversely, worker *i* will invest 1 − *x*_*i*_ effort into task *F*. This approach is closely related to the familiar concept of a response probability: Under the assumption that there is a response probability *p*_*i*_ for worker *i* to engage with task *T* when faced with the choice between *F* and *T*, the expected amount of effort invested in *T* is directly proportional to *p*_*i*_ (and thus to *x*_*i*_).

The state of the colony at any time is given by a vector (*x*_*i*_)_*i*=1…*N*_. We assume that workers interactions are restricted to groups of size *n*, where *n < N*. This physical and spatial constraints. For example, a fanning worker will cool brood only locally, not in every location of the brood chamber. Likewise, social interactions can only take place when workers are in proximity and are thus limited to smaller groups at any point of time.

Individual payoffs depend on *x*_*i*_, as well as on the collective effort invested by all workers in the group *X* = {*x*_*j*_|*j* = 1, 2, …, *n*}. More specifically, worker *i* receives payoff

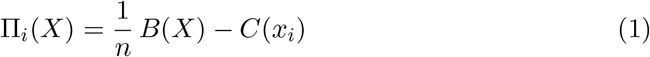

where *B*(*X*) is the total benefit for the group, and *C*(*x*_*i*_) is the cost for worker *i*. This reflects that benefits arising from both tasks are shared, whereas costs are borne individually.

Ensuring colony fitness requires performing both tasks. In the context of our protypical tasks, poor maintenance of nest temperature can slow down the development of the brood and some brood may not survive if there is a shortage of food intake. Hence, the total benefit is multiplicative in the benefits of each task.

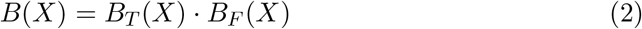

where *B*_*T*_(*X*), *B*_*F*_(*X*) are the benefits of task *T* and *F*, respectively.

Costs, on the other hand, are additive:

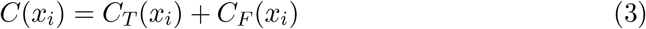

where *C*_*T*_(*x*_*i*_), *C*_*F*_(*x*_*i*_) are the costs of task *T* and *F*, respectively.

The cost of the regulatory task *C*_*T*_(*x*_*i*_) is linear in the effort *x*_*i*_. Consider cooling fanning in bees: the amount of energy required depends on physiological factors of the individual, but it is proportionate to the length and intensity of the fanning activity (effort). For the maximising task we assume marginally decreasing costs, i.e., 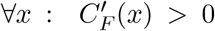 and 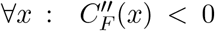. This reflects efficiency improvement through task experience. A foraging bee may become more efficient at finding high value flowers and thus the marginal investment for an additional unit of food decreases.

To investigate how colonies perform in different environmental conditions we introduce two further parameters *b* and *r* that link cost and benefit to the characteristics of the environment (Figure 1). Parameter *b* captures the ratio between the benefit and cost of task *F*. Larger values of *b* represent abundant ecologies where foraging is cheap, whereas small values of *b* indicate that foraging is more costly. Similarly, *r* is the cost ratio of *T* and *F*, i.e., *r*< 1 implies *F* is more expensive than *T* per unit effort, and vice versa. Full details are given in Section A, S1 Appendix.

We now turn to defining how individuals may use their payoffs to adjust their trait values. We consider and simulate two widespread update rules that are at two opposite ends of the spectrum of individual information processing: *individual learning* and *social learning*. We implement this process in an agent-based model and compare the outcomes to an analytic treatment using adaptive dynamics [16].

The simulations start from a homogeneous population. At each time-step, individuals form random groups of size *n*, and play the game described in Equation 1, receiving a payoff according to the collective investment in the task and the individual costs. The population then adjusts strategies based on the learning mechanism.

**Figure 1.**
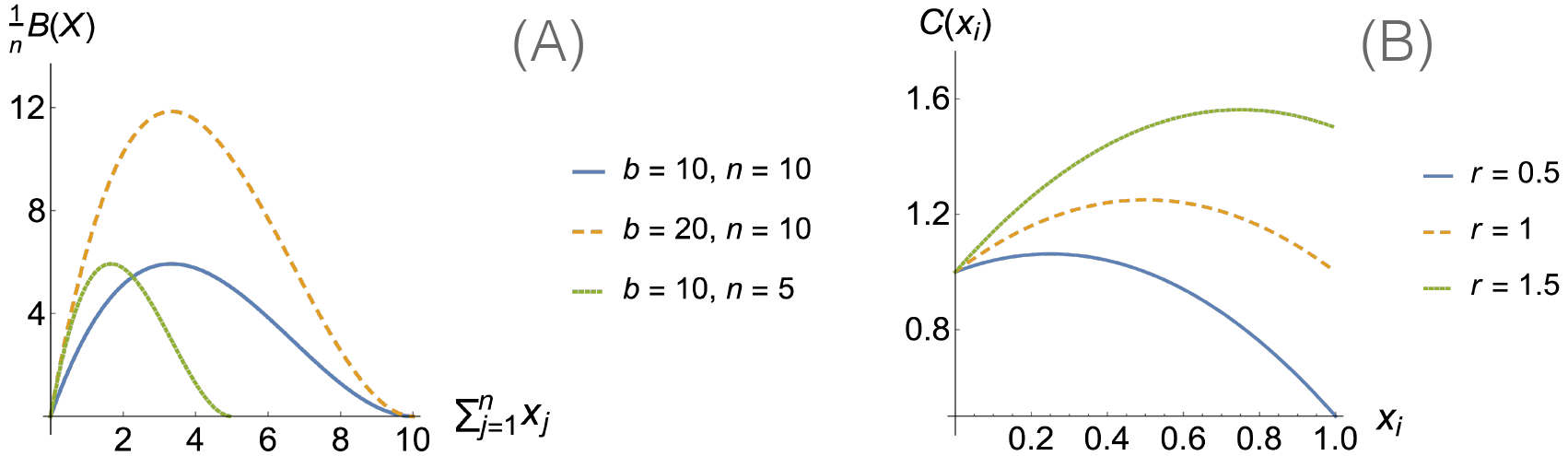
Examples benefit and cost functions. **(A):** Individual benefit as a function of collective group effort. The benefit reaches a maximal value only if the proportion of group workforce allocated to both tasks is appropriate. **(B)**: Individual cost as a function of her own strategy. For *r* < 1, foraging tends to be more expensive than regulation and the cost is minimised for *x*_*i*_ = 1. For *r* > 1, regulation tends to be more expensive than foraging and the cost is minimised for *x*_*i*_ = 0. For *r* = 1 both tasks tend to be equally costly.

### Individual learning

Individual learning is likely to influence the task performance and responsiveness of colony members in social insects [47, 66, 50, 14]. Individuals can adapt their strategies by exploring the current context with previously acquired information [67, 36]. Here we use a simple assumption that individuals assess and improve their strategies by making comparisons between their current and previous task performance. More specifically, each individual explores a new strategy with a small probability, and switches to it only if the new variant provides a larger payoff. This is akin to the ideas of reinforcement learning [42, 71] and stochastic hill-climbing [60].

We run an agent based simulation describing how the population changes across discrete time-steps. A heterogenous population is given by a set of *x*_*i*_ values, for *i* = 1…*N*. In each learning period, each individual will explore a new strategy with probability *β*. If individual *i* is chosen to explore a new strategy, her new trait is sampled from a normal distribution with mean *x*_*i*_, and variance *γ* – the later can be conceived as an exploration parameter. Individual *i* will adopt her new strategy only if it outperforms the current strategy across *k* games. The process is described in detail in Algorithm 1.

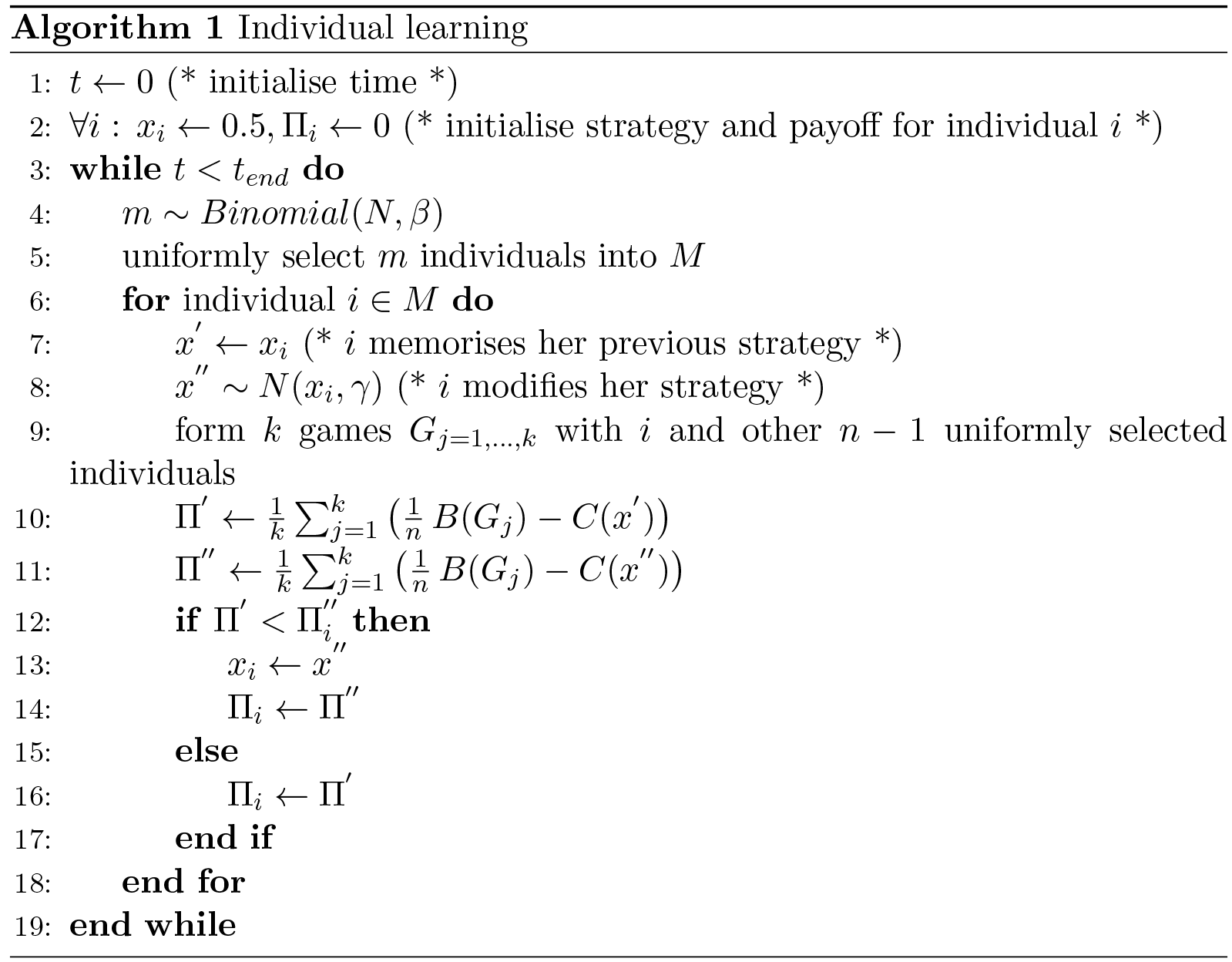

### Social learning

Although not widely discussed in the context of social insects [36], social learning is found by empirical studies in bee foraging [54, 36, 82, 53, 55, 56, 50]. Social learning has also been established in social spiders [64, 65], with the important distinction that these are not eusocial. Due to the potential complexity involved, we make no specific assumptions on the proximate mechanisms of social information exchange. Here we simply assume that individuals are more likely to copy the strategies of others who are successful. Each individual also can explore a new strategy with a small probability. This is similar to the Wright-Fisher process [41] or Roulette-wheel selection in Evolutionary Computation [23].

We run an agent-based simulation describing how the population changes across discrete time-steps. Similar to the case of individual learning, a heterogenous population is given by a set of *x*_*i*_ values, for *i* = 1…*N*. Likewise, during each learning period each individual will explore a new strategy with probability *β*. If individual *i* is chosen to explore a new strategy, her new trait is sampled from a normal distribution with mean *x*_*i*_, and variance *γ*. In contrast to the case of individual learning, here individual *i* will adopt her new strategy by copying successful strategies in the population with a higher probability. The process is described in detail in Algorithm 2.

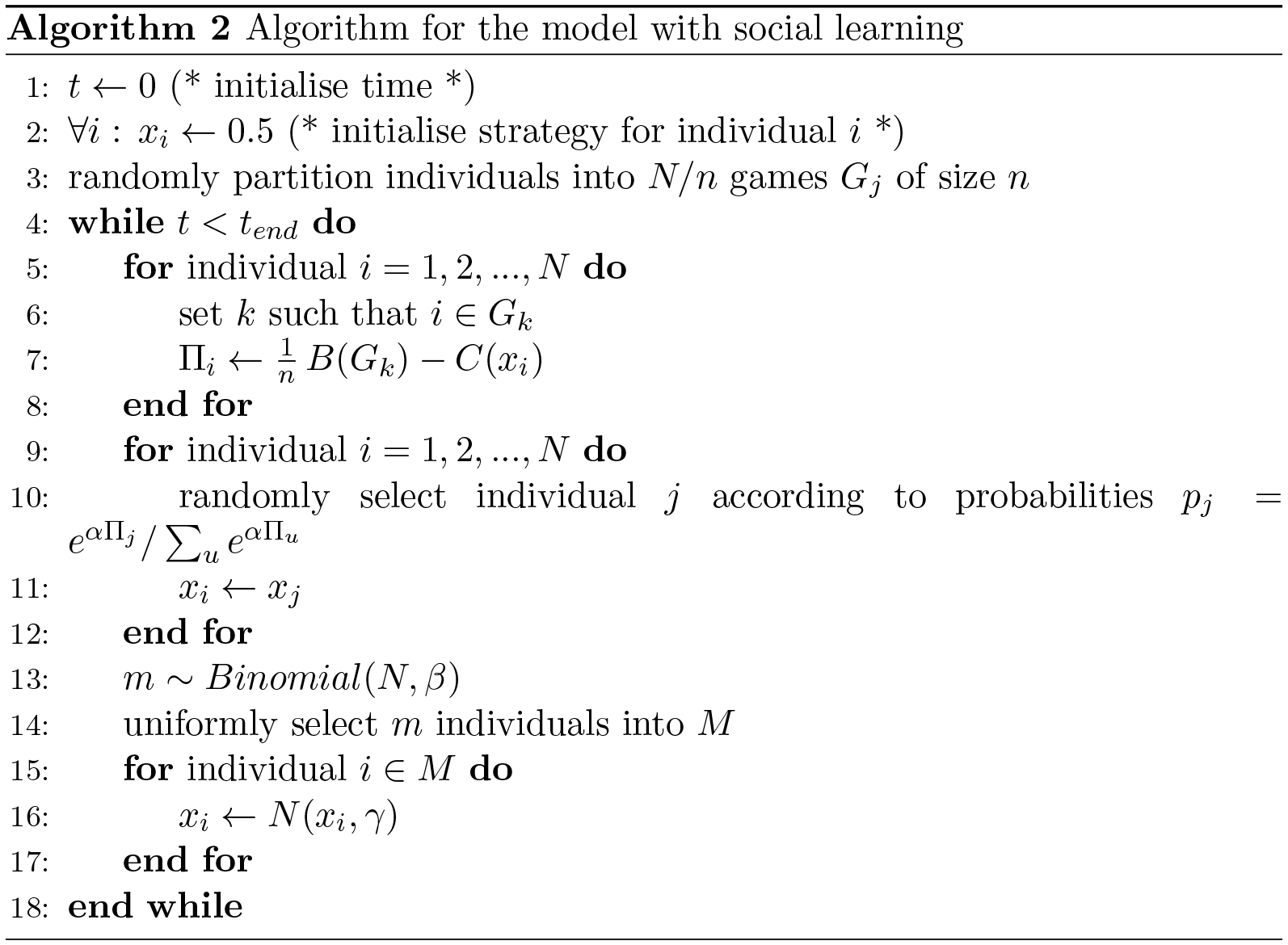

## Results

### Long-term dynamics

Our results show that colony-level task specialisation can emerge from the interaction dynamics between individuals and their environments alone (see strong specialisation in Figure 2 and Figure 3). Under a certain range of environment conditions, the workforce of colonies initially consisting of individuals with identical strategies split into different groups. In each group individuals specialise into a single task (*strong specialisation*), which is driven by social interactions.

We also find that different environmental conditions (characterised by *b* and *r*) can cause variation of task allocation patterns, even in the absence of variation in the underlying individual-based mechanisms. As shown in Figure 2 and Figure 3, strong specialisation tends to emerge in environments with scarce or poor food resources (when *b* is small). As the quality of food resources in the environment improves (*b* increases), individuals are less likely to strongly specialise. Individuals may still prefer one task over the other (*x*_*i*_ ≠ 1/2) to balance global demands, but their strategies tend to be consistent across the colony (*weak specialisation*).

**Figure 2.**
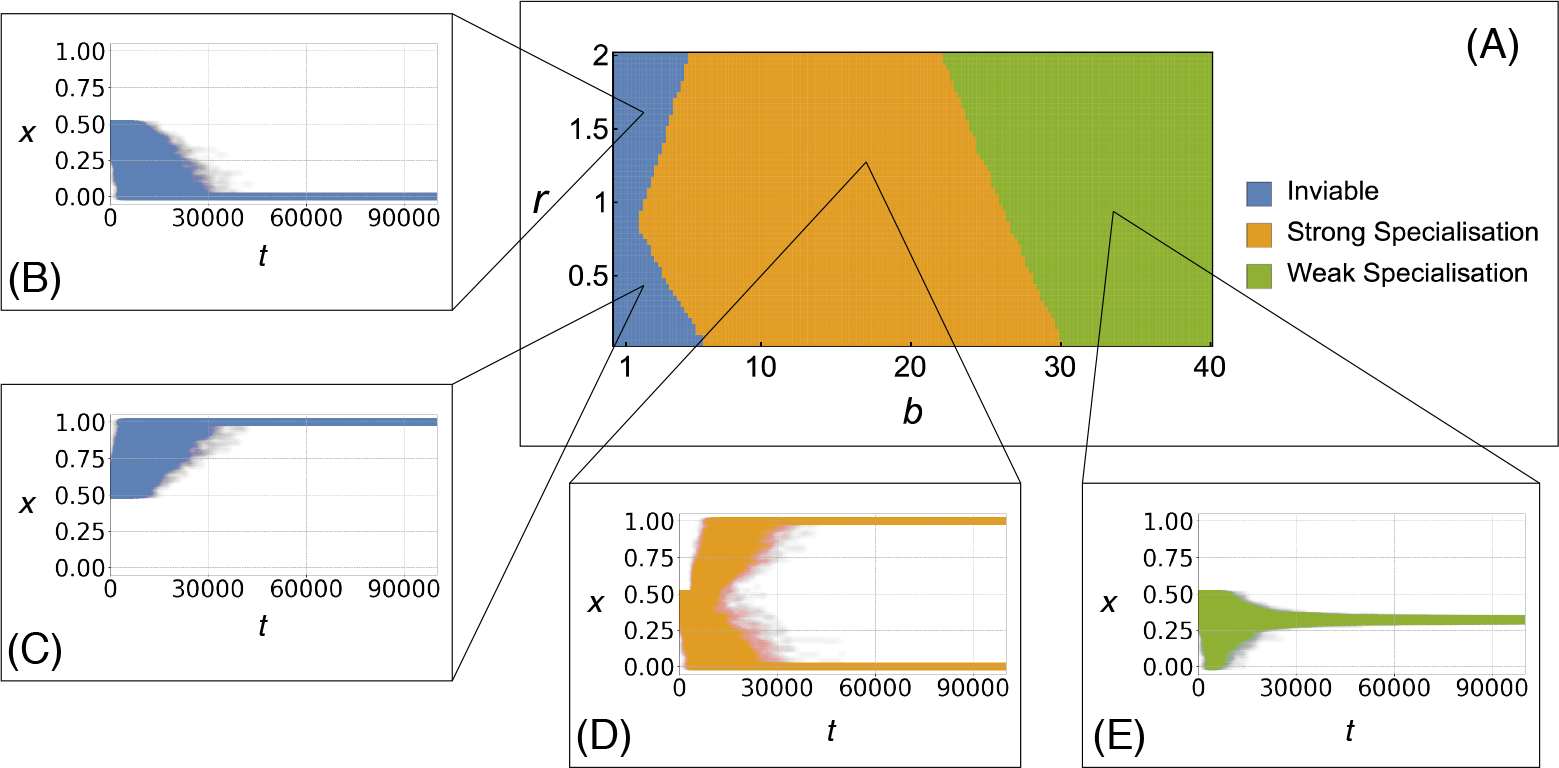
Behavioural patterns for individual learning. **(A):** The phase space diagram describes the long-term outcomes of task allocation for the model with individual learning over a range of values for parameters *b* and *r*. The inset figures **(B)**, **(C)**, **(D)** and **(E)** are scatter plots of trait values for all individuals against time. They show three typical cases of how individual strategies (*x*) in a colony develop over time (*t*). A colony is inviable (**(B)** and **(C)**) if the average payoff of individuals is not positive, which means that they fail to coordinate and only respond to a single task. Strong specialisation **(D)** means that the workforce of a colony splits into two different groups each of which focuses exclusively on a single task. Weak specialisation **(E)** means that each individual invests her effort on both tasks. Section C, S1 Appendix describes the procedure we follow to classify these results from our simulations.

The dynamics of our task-allocation games are similar to those arising from the continuous Snowdrift game [17]. They can be analysed using adaptive dynamics [25]. A full analytical treatment that confirms our simulation results for individual learning and social learning is given in Section B.1, S1 Appendix. This theoretical prediction, shown in Figure 4, matches our simulation results presented in Figure 2(A) and Figure 3(A).

### Efficiency analysis

Although colony optimality is not the driver of the emerging colony organisation, we can use it to quantify group-level efficiency. To do so we use the notion of *relative colony performance*, which is the ratio between the average payoff achieved by individuals in a colony and the level of payoff that could be achieved with optimal workforce allocation. This concept is similar to the *price of anarchy*, used in Computer Science to quantify the cost of decentralised organisation [52].

**Figure 3.**
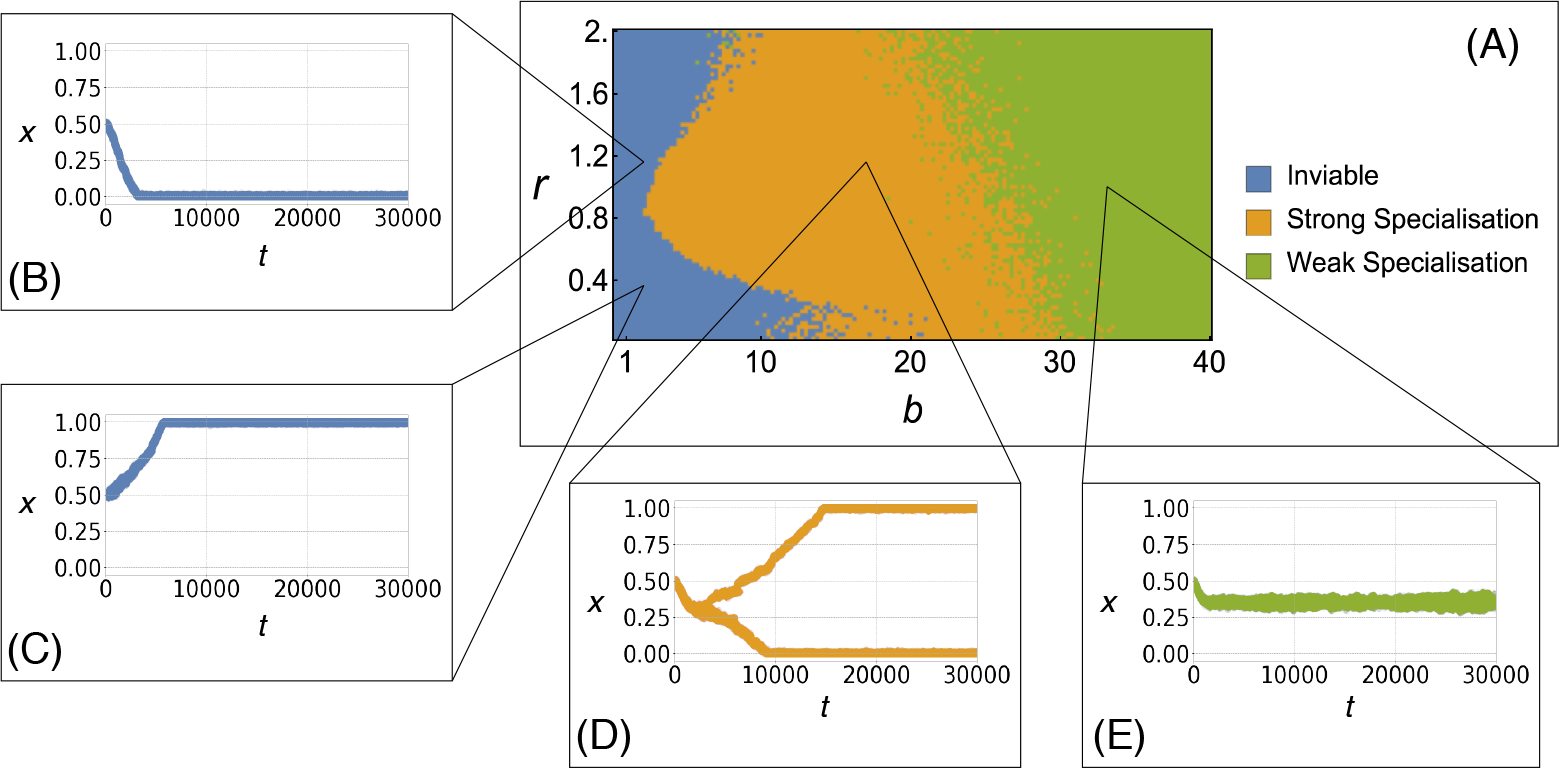
Behavioural patterns for social learning. **(A):** The phase space diagram gives the long-term patterns of task allocation of colonies based on the model with social learning in a range of values for parameters *b* and *r*. The inset figures **(B)**, **(C)**, **(D)**and **(E)** are scatter plots of trait values for all individuals against time. They show three typical cases of how individual strategies (*x*) in a colony develop over time (*t*). A colony can be either inviable (**(B)** and **(C)**), strongly specialised **(D)** or weakly specialised **(E)**. Being inviable means that the average payoff of individuals in a colony is not positive and the overall task allocation is out of balance as individuals only respond to a single task. Strong specialisation means that the workforce of a colony splits in two groups each of which specialise in a single task. Weak specialisation means that each individual spends her energy on both tasks. Section C, S1 Appendix describes the procedure we use to classify the above results from the simulations.

To measure the colony performance we take parameters *b* and *c* and determine if the long term outcome is a monomorphous or dimorphous population, by computing the equilibrium *x*\* and inspecting the higher order derivatives of the invasion fitness at that point. If the dynamics is monomorphous, the average payoff in equilibrium can be calculated directly using *b*, *c* and *x*\*, because all the individuals will choose strategy *x*\*. If the population is dimorphic, we use the replicator equation to determine the frequencies at which specialists choose to fully engage in either task, *ρ*\*. With *ρ*\*, *b* and *c* we can compute the average payoff in equilibrium. We divide the average payoff by the optimal payoff, derived from optimising the group payoff. Details for the calculations are given in Section B, S1 Appendix.

**Figure 4.**
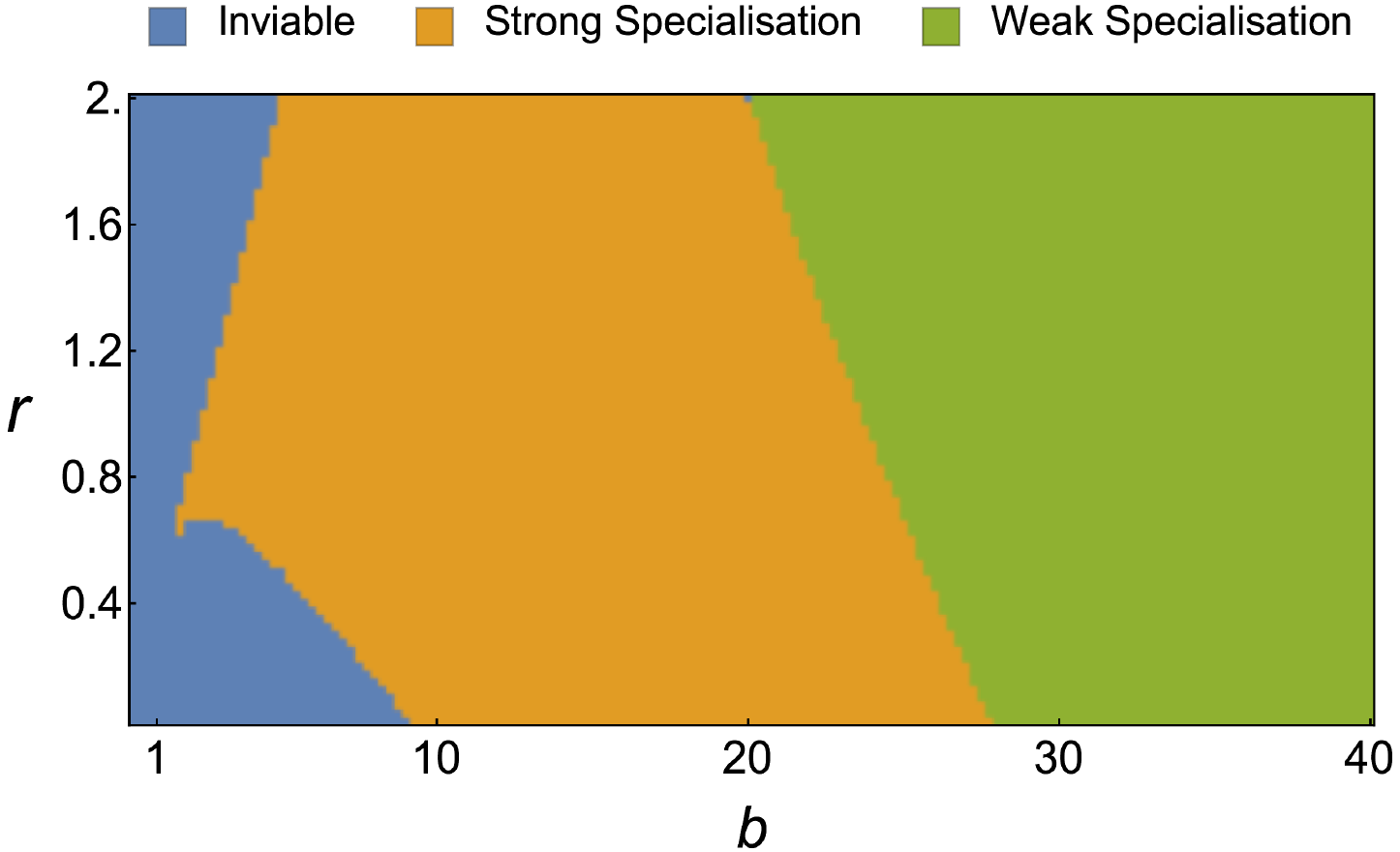
Behavioural patterns predicted by adaptive dynamics. The above diagram describes the patterns of task allocation of colonies based on the mathematical framework of adaptive dynamics in a range of values for parameters *b* and *r* (given *n* = 10).

We find that relative colony performance varies with environmental conditions. Different ecologies result in different forms of specialisation, which in turn leads to different levels of relative colony performance. A relative colony performance of 1 indicates optimality. Interestingly, weakly specialised colonies turn out to perform close to optimal, while strongly specialised colonies can perform sub-optimally.

This is because, in strongly specialised colonies, individuals need to select a single task into which they invest their entire effort. As benefits are shared evenly, individuals will adjust their strategies in favour of the task that is less costly as long as the impact of doing so on the shared benefit does not outweigh the effect of reduced cost on the level of the individual.

### Dynamic environments

We find that different learning mechanisms can lead to varying behavioural patterns under environmental fluctuations. Our results suggest that individuals in the colonies based on individual learning tend to flexibly adjust their strategies according to the current environmental conditions. As seen in Figure 6, the colony based on individual learning changes from weak specialisation to strong specialisation when foraging becomes less profitable (*b* decreases) and switches back to weak specialisation once the environmental condition has reverted to the original state. This allows the colony to be near optimal efficiency in spite of the environmental fluctuations.

**Figure 5.**
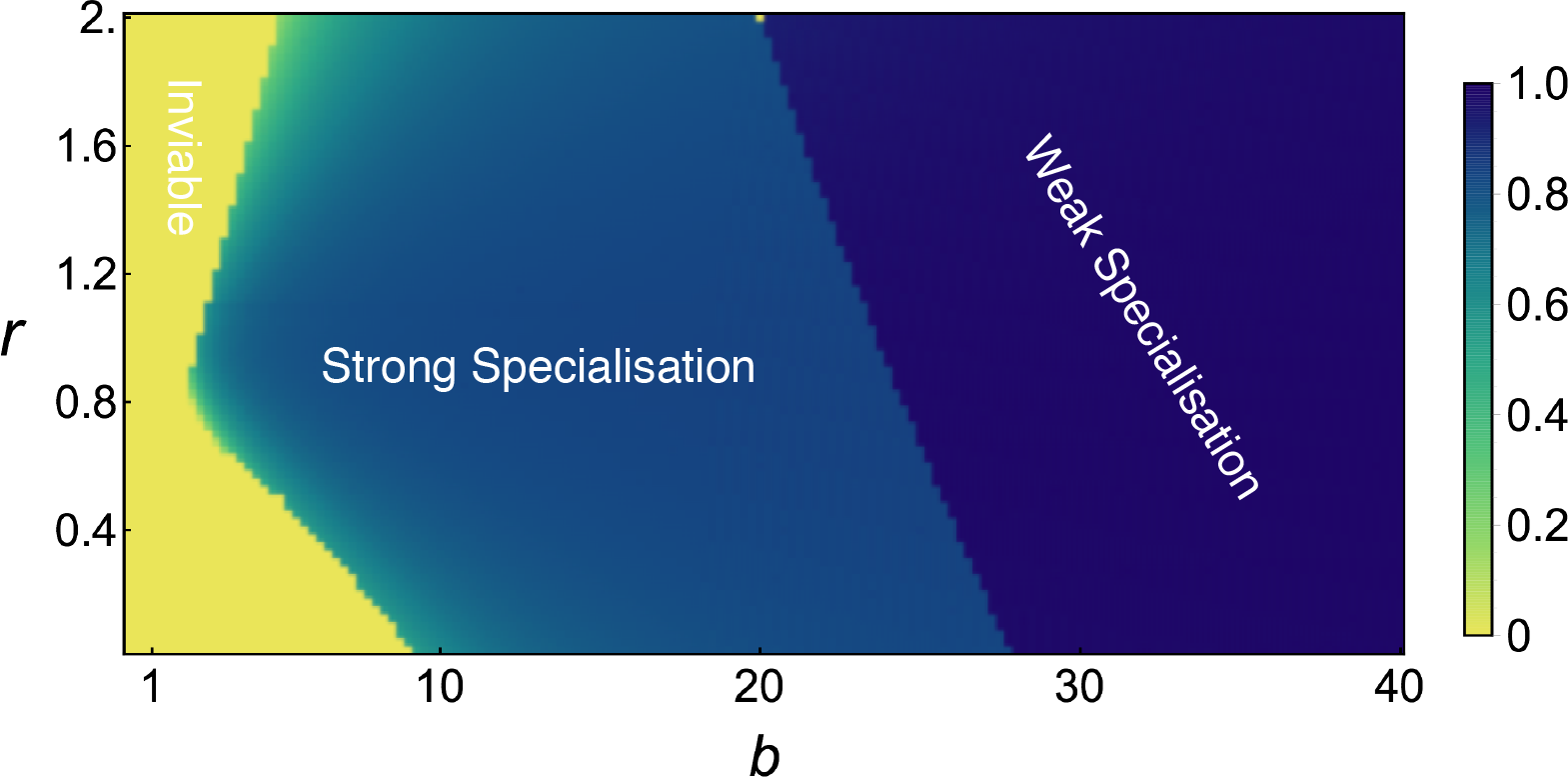
Relative colony performance. For a pair of *b* and *r* values, the efficiency is the ratio between the mean of individual payoffs at a long term and the optimal level that can be theoretically achieved under the environmental condition. Efficiency ranges from 0 to 1. For simplicity, we regard the efficiency of inviable colonies as 0.

Surprisingly, for social learning, our results suggest that the colony-level patterns of task allocation do not only depend on the current condition of the environment, but also on the history of these conditions. As illustrated in Figure 6, when *b* decreases, the colony based on social learning changes from weak specialisation to strong specialisation. However, when the environmental conditions return to the original state, the pattern of task allocation at the colony level does not revert to the original pattern of organisation, and the colony cannot regain its original performance. In other words, suboptimal outcomes arise.

**Figure 6.**
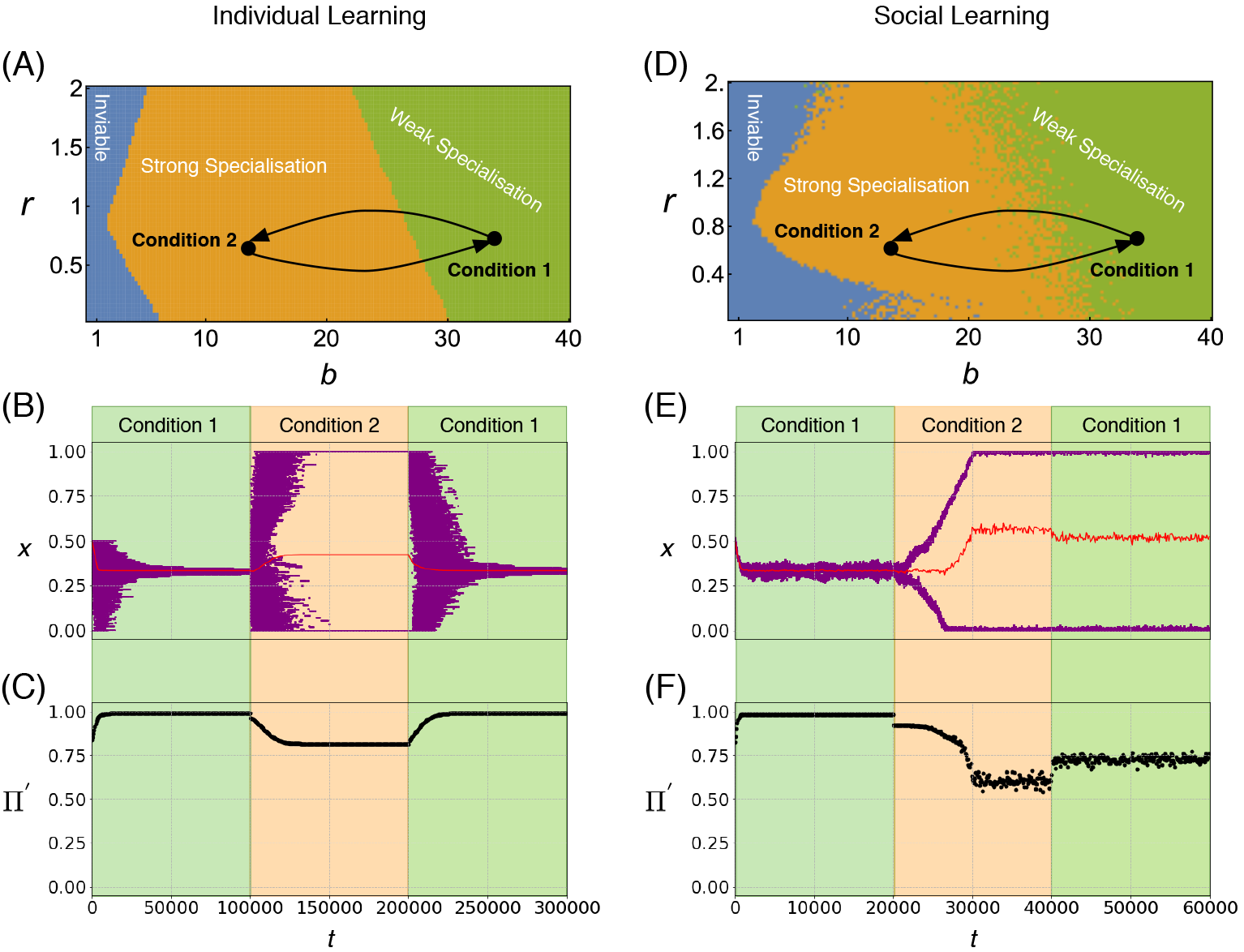
Individual learning and social learning in dynamic environments. We compare the dynamics of learning when the conditions of the environment change between two conditions in which social organisation differs. Panels **(A)**, **(B)** and **(C)** refer to individual learning, while panels **(D)**, **(E)** and **(F)** refer to social learning. For individual learning, **(B)** depicts the dynamics of individual strategies (*x*) and **(C)** shows the efficiency (Π’) in a colony over time (*t*) under environmental fluctuations. The environment is set as three stages: (1) Condition 1 (*b* = 40) for *t* ∈ [0, 100000]; (2) Condition 2 (*b* is switched to 10) from *t* = 100000 to *t* = 200000; (3) Condition 1 (*b* is switched back to 40) until the end of simulation (*N* = 2000, *t*_*end*_ = 300000, *n* = 10, *r* = 0.7, *β* = 0.001, *γ* = 0.1). Similarly, for social learning, **(E)** gives the dynamics of individual strategies with the mean and **(F)** illustrates the relative colony performance under environmental fluctuations. The environment is also set as three stages: (1) Condition 1 (*b* = 40) for *t* ∈ [0, 20000]; (2) Condition 2 (*b* is switched to 10) from *t* = 20000 to *t* = 40000; (3) Condition 1 (*b* is switched back to 40) until the end of simulation (*N* = 2000, *t*_*end*_ = 60000, *n* = 10, *r* = 0.7, *α* = 2.5, *β* = 0.01, *γ* = 0.005). Here **(A)** and **(D)** correspond to the region diagrams in Figure 2 and Figure 3 respectively. **(B)** and **(E)** are scatter plots of trait values for all individuals against time with the mean trait value indicated by a red line.

## Discussion

We introduce a new modelling framework for task allocation in social insects, based on evolutionary game theory, and use this framework to analyse different behavioural patterns in terms of specialisation. This is motivated by the fact that conventional frameworks do not sufficiently address two aspects that are recognised as crucial to the investigation of task allocation: the role of environmental conditions and the role of social interaction [30]. Specifically, we ask two questions: (1) what are the factors that can determine whether specialisation arises and (2) how do different behavioural patterns relate to the overall colony efficiency.

Two most commonly used types of models in task allocation, response-threshold models [3, 47] and foraging-for-work models [79, 78], share the characteristics that groups of independent individuals are described as acting in parallel. Interaction between these individuals only takes place indirectly through modification of the environment (stigmergy). There is no genuine place for social interactions in these models. In contrast, our EGT-based approach integrates social interaction explicitly as a fundamental component.

Furthermore, the EGT framework allows us to easily exchange the underlying mechanisms of learning and compare the effects of such changes. We have exploited this capability to compare the dynamics of social learning with that of individual learning. Our analysis reveals similar outcomes in behavioural patterns of task allocation by comparing individual learning and social learning. In our simulations, both types lead to different types of specialisation depending on the environment (or, in broader terms, the ecology). The most striking aspect of this is that the ecology alone can determine which type of specialisation arises, without any changes in the underlying proximate mechanisms of task selection.

One of the core interests in the study of task allocation is to investigate the primary sources of variation in workers’ task preference and ultimately specialisation [30]. Many studies regard inherent inter-individual differentiation as the main cause [47]. Some studies have shown that specialisation can arise in colonies of initially identical workers either from reinforcement via individual experience [77] or from spatially localised task demands [79, 49]. Our results show a new alternative driver for specialisation, namely social interaction between individuals (see Figure 2 and Figure 3).

Our model is based on costs and benefits perceived by the individual. In the context of eusocial insects it is important to point out that this includes the possibility of a proxy for colony level benefit that can be perceived by an individual. On the other hand, in animals that are social but not eusocial, such as social spiders, the benefit accrues directly to the individual.

It is known that experience-based reinforcement is likely to influence workers’ decision-making in task selection [47]. Our model with individual learning predicts that when the resources are less abundant, colonies tend to behave in strong specialisation with different tasks. This is congruent with previous theoretical models of reinforcement of individual experience in task allocation [77]. However, it is interesting to explore whether colony workforce may change into a different form (weak specialisation) if the environmental condition becomes less challenging.

There is clear evidence that social learning matters in social insect colonies [26, 36]. However, little is known about the exact mechanisms of social learning in relation to task allocation, and there is a clear need for further empirical work in this regard. For our models we have assumed one of the most basic forms of social learning. Since there is insufficient knowledge about the details of the real biological mechanisms that may be at work, this provides a reasonable starting point. Importantly, the dynamics that these assumptions lead to, so-called replicator dynamics [38], are qualitatively stable for a reasonably broad range of changes in the detailed learning mechanisms. Replicator dynamics arises in a surprisingly wide range of different learning scenarios [71, 42]. It thus provides a good basis for a hypothetical discussion in the absence of more precise knowledge.

We were able to show that the qualitative behaviour remains unchanged when switching between two extreme ends of the spectrum of learning mechanisms: individual and social. This suggests that the finer details of the social learning mechanisms that may actually be at play will only have a limited impact on this qualitative behaviour.

Our results suggest that individual learning leads to higher colony performance under fluctuating environmental conditions. One may thus expect that social learning is selected against in environment types where it does not achieve high efficiency. However, social learning may arguably provide other benefits to the colony, most importantly a mechanism to spread information through the colony in the absence of local cues. Empirical studies suggest that workers can recognise the tasks that others perform simply by chemical cues or antennal contact [27, 31]. Spatial movement is widely observed and is likely to influence task allocation in social insects [28, 9, 73, 48, 8]. It is thus conceivable that these benefits outweigh the potential price that is paid in terms of overall performance.

In environment types where both weak and strong specialisation can exist, weak specialisation can be more efficient than strong specialisation (see social learning in Figure 6). This seems to contradict an often made assumption that strong specialisation is one of the ways how colonies achieve higher efficiency [62, 10]. However, it is established that this is not always the case and that the evidence for this is not consistent [14]. Our models show strong specialisation may, in some circumstances, be an emergent byproduct of the proximate mechanisms that determine individual task selection rather than a fitness improving outcome (and thus directly selected for).

However, strong specialisation in a social learning context may convey a fitness benefit in fluctuating environments. To adjust to changing conditions, weakly specialised colonies must depend on individual-based exploration of the strategy space by a small proportion of workers. In contrast, strongly-specialised colonies can benefit from imitation between different specialised group to adjust the ratio of workers. In the dynamics of game theory, the process of exploring new strategies is generally assumed to occur on a much longer timescale than strategy imitation [6]. It is thus not unreasonable to expect that strongly specialised colonies may adapt faster and more flexibly to changing conditions.

Our task-allocation game is similar to a continuous Snowdrift game [17], in which the benefit is shared by all individuals at the group level, but the costs are strategy-dependent at the individual level and tend to be different across individuals. Both games can be used to explore features of cooperation and illustrate a principle called “Tragedy of the Commune” [17]. This principle describes non-uniform investment across a group that receives uniform benefit: some individuals significantly contribute to generating a common good, while some “free loaders” invest less or nothing and still reap the same benefits. Ultimately, this may give us a new perspective to tackle the puzzle of “lazy” workers in social insect colonies [12, 10, 11, 37, 13].

There are some factors that may influence the outcomes of the modelled process and whose influence remains to be investigated in more detail. One of these is the “interaction range” of individuals. It is well known that individuals in a colony, due to their physical or spatial limitations, typically sample and respond to localised cues as a proxy for the global situation [27]. In our model this is reflected by letting individuals interact in smaller subgroups (the games), to which their information gathering is limited at any point of time. The size of a game then is a proxy for the scale of interaction in the colony. Game size is a factor that can influence the ultimate outcomes of the EGT models [5], and the impact of games size on our models remains to be investigated. The strength of our modelling approach is that it gives us a principled way to investigate the influence of such factors.

As a starting point we have modelled the allocation between a homeostatic task and a maximising task. While this addresses some fundamental aspects, the range of possible task types is obviously much larger. Different cost and benefit functions, associated with different task types, must be expected to have a significant influence on the outcomes of the models. Game theory research has shown that a qualitative classification of games can go a long way in determining long-term behaviour [1]. This allows us to abstract from the exact quantitative nature of these functions and to switch to a qualitative perspective. This is a powerful concept, since the exact qualitative nature of cost and benefit functions can usually not be ascertained. The hope is that switching to a qualitative view will allow us to focus the discussion on different fundamental types of tasks that are competing for attention. This should impact both, theoretical work as well as experiments. This approach has the potential to address central questions in social insect task allocation.

## Appendix A Details of the payoff function

As task *T* needs to be controlled at a certain level, under or over performing task *T* can reduce *B*_*T*_ (*X*). We use a simple way to model

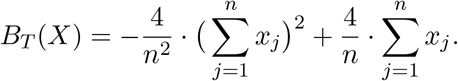

Here *B*_*T*_ (*X*) is assumed to achieve the maximum value, which is normalised between 0 and 1, when half of the workforce in the game is engaged in task *T* and to be 0 when none or all of workers in the game are engaged. As task *F* is a maximising task, which implies, for example, the more food is collected, the more brood can survive, *B*_*F*_ (*X*) is simply assumed to be linear

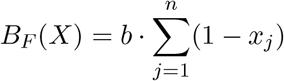

where *b* is coefficient ratio between the benefit of task *F* and the cost of task *F*.

We assume the cost of a homeostatic task to be linear in individual effort and thus define

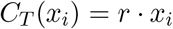

where *r* is coefficient ratio of the costs between task *T* and task *F*. We assume that *C*_*F*_ (*x*_*i*_) is marginally decreasing with the effort in task *F*, indicating the scenarios in which foragers initially need to spend more effort exploring their neighbourhood and once they become familiar with the surrounding areas of food resources, the cost for them tend to be less than the initial stage. As a result, we simply assume

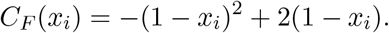

Here the cost of task *F* for a worker who engage fully in task *F* per time-period is assumed to be 1 unit.

## Appendix B Theoretical analysis

## B.1 Analysis for a monomorphous population: adaptive dynamics

## B.1.1 Equilibrium points

We follow the technique used by [17] to study non-linear public good games in continuous traits. In a game of size *n*, with *n*−1 type-I workers of strategy *x* and 1 type-II worker of strategy *y* (*x*, *y* ∈ [0, 1]), the growth rate (invasion fitness) of the type-II worker is

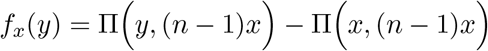

Where

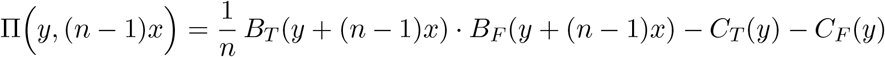

and

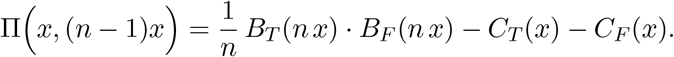

Thus, the selection gradient is

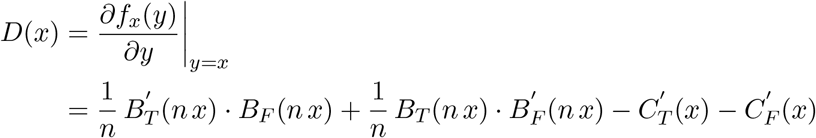

Then the singular strategies are given as solutions of

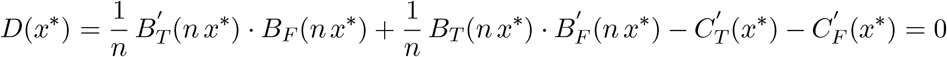

Both strong specialisation and weak specialisation require the condition that there exists such *x*\* ∈ [0, 1] that *x*\* is convergency stable

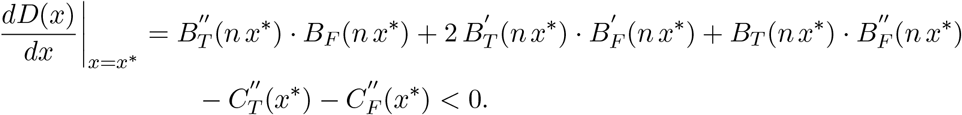

## B.1.2 Branching condition (weak or strong specialisation)

In addition to the above condition, strong specialisation emerges if

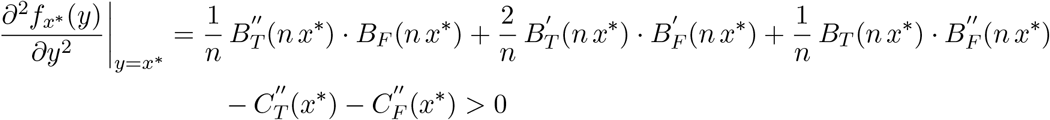

and weak specialisation requires

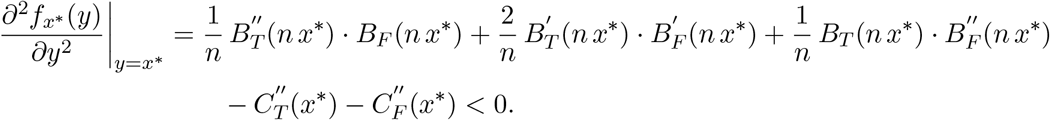

In other cases such as when *x*\* is convergency unstable, colonies tend to become inviable based on our payoff function, as one task out of the two is abandoned.

## B.2 Analysis of dimorphic populations: replicator equation

We provide mathematical analysis of our task-allocation game for the case of strong specialisation (only two types of strategies involved) based on the replicator equation [76, 72].

In a well-mixed colony of large size, the proportion of type-I individuals of strategy *x* is *ρ* and the proportion of type-II individuals of strategy *y* is 1 − *ρ* (*x*, *y* ∈ [0, 1]).

Given that the size of game is *n*, the dynamics of the fraction of individuals of type I individuals is given by:

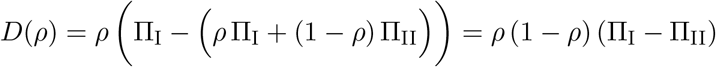

where

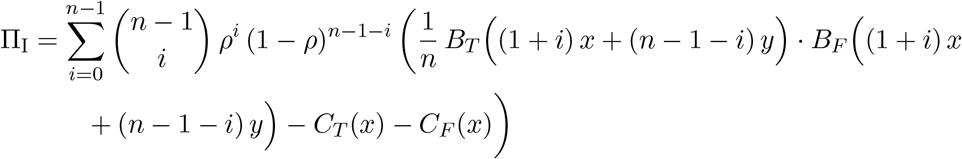

and

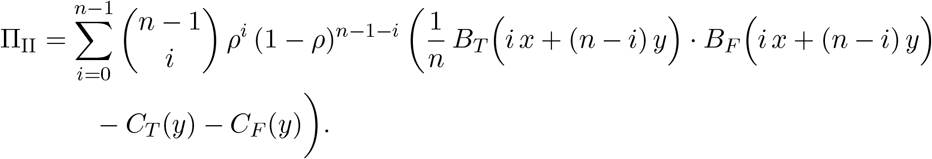

The solutions of *D*(*ρ*\*) = 0 for *ρ*\* ∈ (0, 1) give the stable equilibrium for the long-term dynamics of individual strategies in a colony. The quantity *ρ*\* is the proportion of strongly specialised individuals who fully engage in foraging.

## B.3 Optimal payoff

To evaluate the efficiency achieved by the colony we also need to know the optimal level associated with different environmental conditions (illustrated in Figure S7). To find this, for each pair of *b* and *r*, we optimise the mean of individual payoffs with potentially different strategies in a game of size *n* using Differential Evolution [75], a stochastic population-based heuristic method for global optimisation (implemented by *differential_evolution* in the package *optimize* of Scipy, Version 0.19.0).

**Figure S7.**
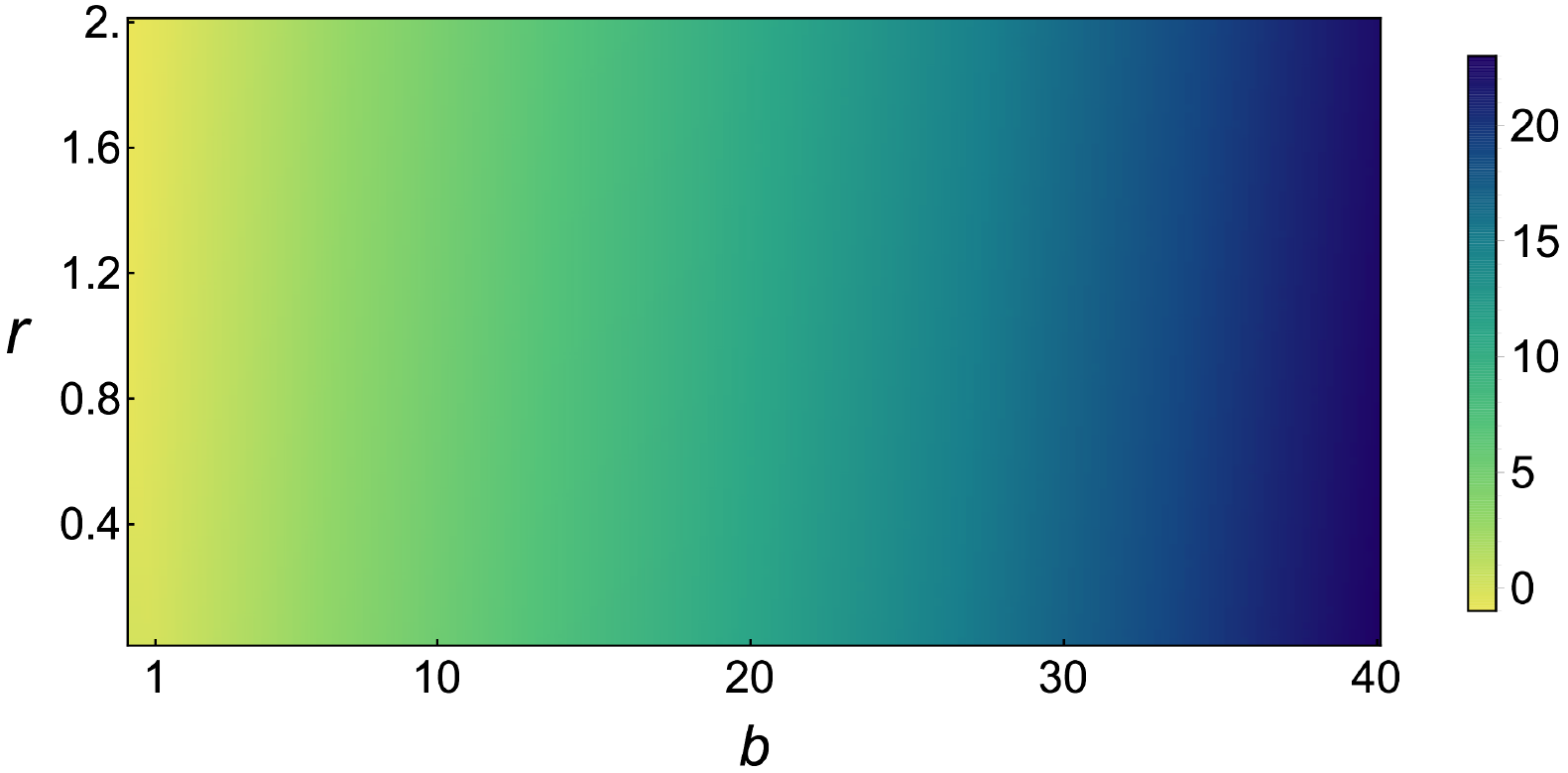
Optimal individual payoff. This diagram gives the optimal payoff that an individual can achieve in a game (*n* = 10) under in a range of values for parameters *b* and *r*.

## Appendix C Classification of simulation results

For the models with individual learning and social learning, the colonies with the non-positive mean of workers’ payoffs are classified under being inviable (for details of the mean of individual payoffs, see Figure S8(a) and Figure S9(a)). The other colonies are tentatively regarded under strong specialisation if the standard deviation of workers’ strategies exceeds a certain level (set as 0.1) or weak specialisation otherwise (for details of the standard deviation of individual strategies, see Figure S8(b) and Figure S9(b)).

**Figure S8.**
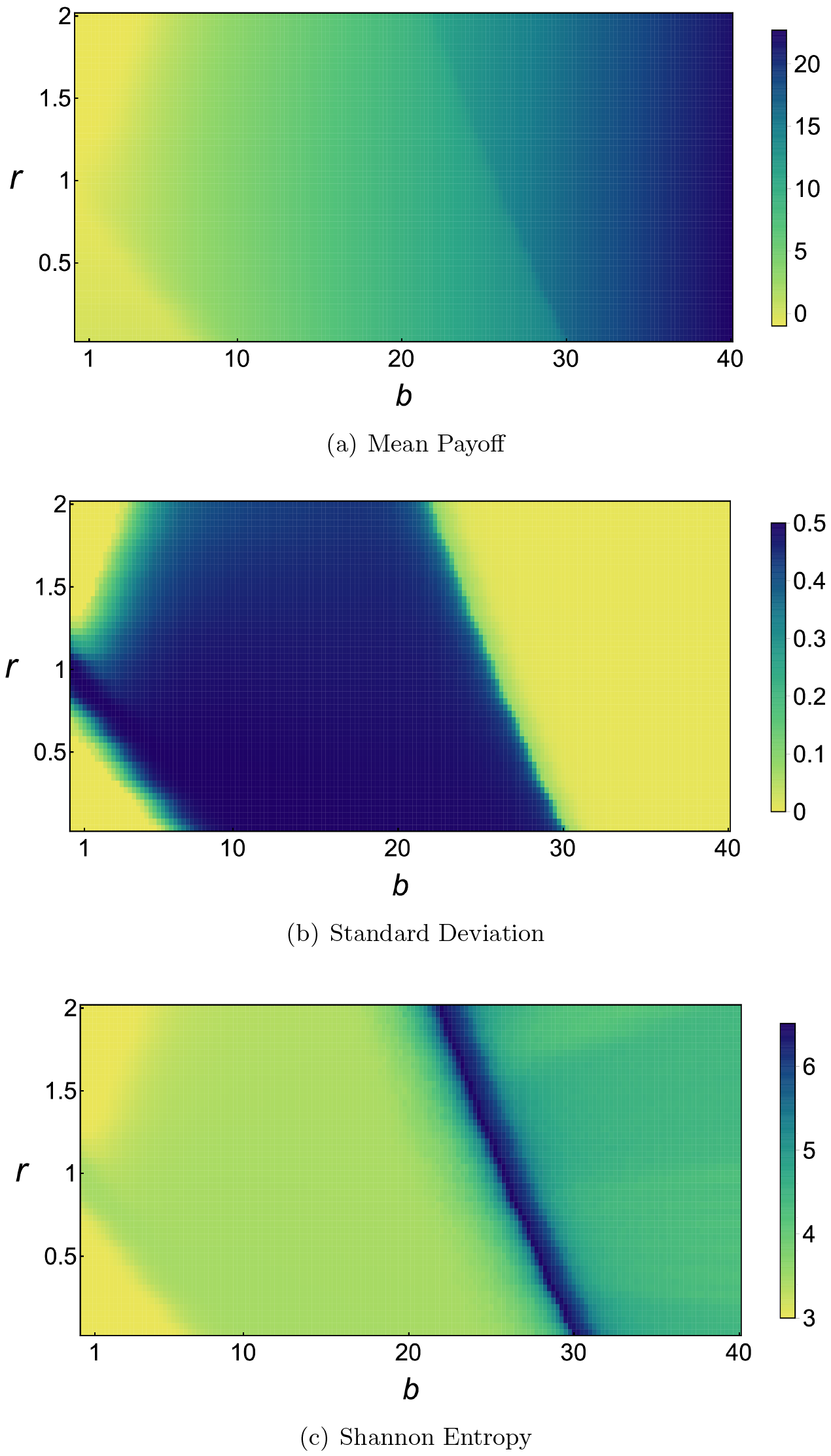
Results of the model with individual learning (*N* = 2000, *t*_*end*_ = 100000, *n* = 10, *β* = 0.001, *γ* = 0.1). In order to highlight the variation within this model under different environmental conditions, each of the above figures has a unique colour scheme.

**Figure S9.**
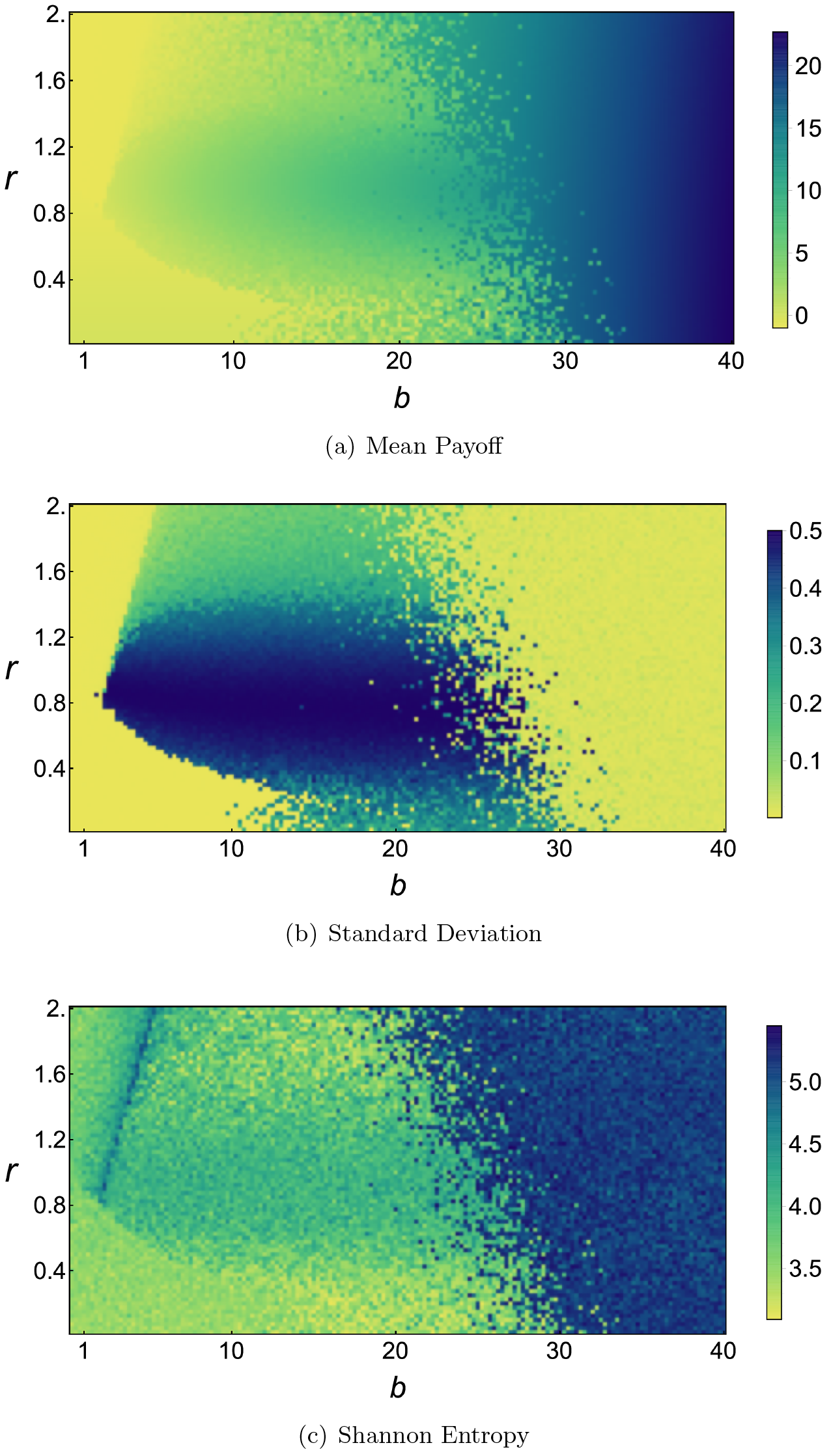
Results of the model with social learning (*N* = 2000, *t*_*end*_ = 30000, *n* = 10, *α* = 2.5, *β* = 0.01, *γ* = 0.005). In order to highlight the variation within this model under different environmental conditions, each of the above figures has a unique colour scheme.

However, a large standard deviation of individual strategies cannot guarantee strong specialisation, as a colony with a wide span of strategies may belong to weak specialisation and correspond to a large standard deviation as well. To capture the span of individual strategies, we verify the above temporary region classification by the Shannon entropy (for details, see Figure S8(c) and Figure S9(c)). For both models, the entropies of individual strategies in colonies with large standard deviation are smaller than those with small standard deviation, which in turn confirms our temporary region classifications. In order to highlight the variation under different environmental conditions, each of the three figures have a unique colour scheme for both individual learning and social learning.

## Appendix D Simulation source code

The source code for our simulations of individual learning and social learning in both static and dynamic environments can be found at https://github.com/rche29/model_task_allocation.

